# Neural population dynamics reveal that motor-targeted intraspinal microstimulation preferentially depresses nociceptive transmission in spinal cord injury-related neuropathic pain

**DOI:** 10.1101/2023.07.27.550880

**Authors:** Jacob G. McPherson, Maria F. Bandres

## Abstract

The purpose of this study is to determine whether intraspinal microstimulation (ISMS) intended to enhance voluntary motor output after spinal cord injury (SCI) modulates neural population-level spinal responsiveness to nociceptive sensory feedback. The study was conducted *in vivo* in three cohorts of rats: neurologically intact, chronic SCI without behavioral signs of neuropathic pain, and chronic SCI with SCI-related neuropathic pain (SCI-NP). Nociceptive sensory feedback was induced by application of graded mechanical pressure to the plantar surface of the hindpaw before, during, and after periods of sub-motor threshold ISMS delivered within the motor pools of the L5 spinal segment. Neural population-level responsiveness to nociceptive feedback was recorded throughout the dorso-ventral extent of the L5 spinal segment using dense multi-channel microelectrode arrays. Whereas motor-targeted ISMS reduced nociceptive transmission across electrodes in neurologically intact animals both during and following stimulation, it was not associated with altered nociceptive transmission in rats with SCI that lacked behavioral signs of neuropathic pain. Surprisingly, nociceptive transmission was reduced both during and following motor-targeted ISMS in rats with SCI-NP, and to an extent comparable to that of neurologically intact animals. The mechanisms underlying the differential anti-nociceptive effects of motor-targeted ISMS are unclear, although they may be related to differences in the intrinsic active membrane properties of spinal neurons across the cohorts. Nevertheless, the results of this study support the notion that it may be possible to purposefully engineer spinal stimulation-based therapies that afford multi-modal rehabilitation benefits, and specifically that it may be possible to do so for the individuals most in need – i.e., those with SCI-related movement impairments and SCI-NP.

## INTRODUCTION

Electrical stimulation of the spinal cord can enhance rehabilitation of function in people living with spinal cord injury. Development of spinal stimulation paradigms to enable voluntary motor output by enhancing the therapeutic benefits of physical rehabilitation is a particularly active area of research. Yet, restoration of movement is far from the only rehabilitation goal for which spinal stimulation may be especially efficacious. SCI-related neuropathic pain (SCI-NP), bowel, bladder and sexual dysfunction, and autonomic dysreflexia are also amongst the top rehabilitation priorities for people living with SCI, and all can be targeted with spinal stimulation (Anderson, 2004; Center, 2021; Faleiros et al., 2023; Snoek et al., 2004).

An emerging class of spinal stimulation-based therapy seeks to afford multi-modal rehabilitation benefits (Bandres et al., 2023b). One approach to developing these therapies is to exploit the off-target effects of spinal stimulation paradigms that were nominally intended to address a single consequence of SCI. In effect, this approach leverages the dense interconnectivity of spinal neurons to purposefully modulate multiple functional networks simultaneously. Towards development of such a therapy, it has recently been demonstrated in pre-clinical rodent models that intraspinal microstimulation (ISMS) intended to enhance voluntary motor output following SCI simultaneously reduces spinal responsiveness to nociceptive transmission, including in rats with chronic SCI-NP (Bandres et al., 2023a; Bandres et al., 2023b; Bandres et al., 2022). This unexpected depressive effect was observed in individual nociceptive specific and wide dynamic range neurons *in vivo*, and manifest as immediate and persistent decreases in discharge rates in response to nociceptive sensory feedback during and following ISMS.

Here, we address a key question left unresolved by these studies: whether neural population-level spinal responsiveness to nociceptive sensory feedback is also decreased by motor-targeted ISMS. Indeed, only a fraction of discrete neurons within a network can be discriminated from multi-unit patterns of neural transmission and reliably tracked during prolonged periods of stimulation and sensory feedback. These limitations render it difficult to infer whether the firing dynamics of reduced samples of well-isolated neurons are reflective of the emergent dynamics of the overall pool. Establishing the extent to which the modulatory actions of ISMS are internally consistent across different scales of neural transmission will yield insights into the potential mechanisms underlying the anti-nociceptive effects of motor-targeted ISMS reported previously, which may in turn facilitate optimization of ISMS paradigms specifically intended to capitalize on this unique multi-modal effect.

## MATERIAL AND METHODS

All procedures were approved by the Institutional Animal Care and Usage Committee of Washington University in St. Louis. The study included 14 adult male Sprague-Dawley rats (∼400-550 g) with moderate to severe sensorimotor deficits following a midline spinal contusion injury at the T8/T9 vertebral border (**Fig. 1A**; Infinite Horizon Impactor, IH-04000; Precision Systems and Instrumentation, LLC; 200 Kilodynes, 0 sec dwell time) and 6 neurologically intact rats. Of the rats with chronic SCI, 7 rats had moderate to severe bilateral hindlimb paralysis and no behavioral signs of SCI-NP and 7 rats had moderate to severe bilateral hindlimb paralysis *and* behavioral signs of SCI-NP.

**Figure 1.**
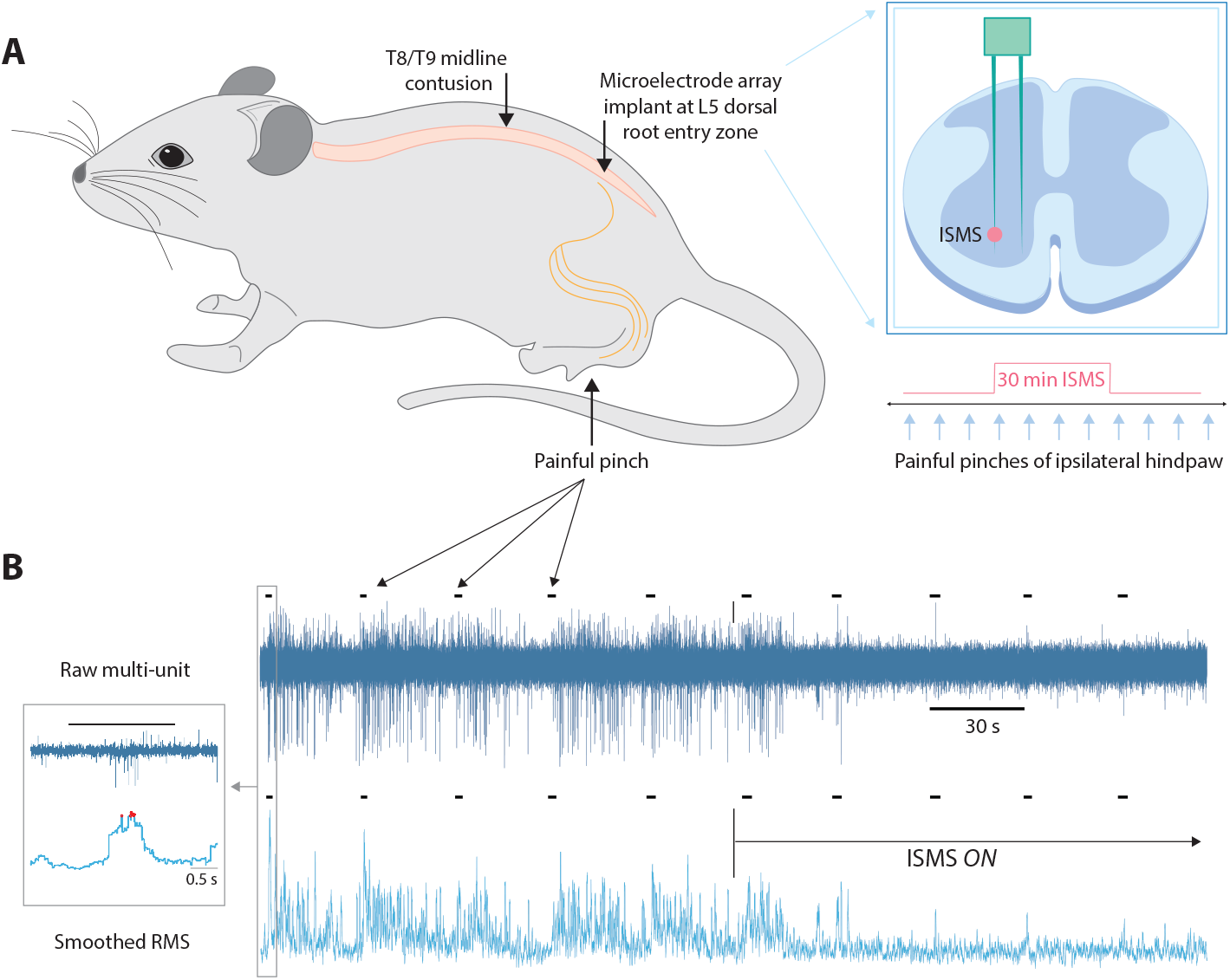
Experimental overview. (**A**) Schematic depictions of: SCI and microelectrode array implant location; microelectrode array placement and dimensions relative to the L5 spinal segment; and general trial structure. (**B**) Representative data illustrating the process and metrics used to quantify nociceptive responsiveness on a given electrode channel. Top trace: raw, multi-unit neural data from one electrode of a microelectrode array; bottom trace: time-locked rectified and smoothed version of the above record; small black hash marks indicate onset and duration of painful pinches of the ipsilateral hindpaw. Data are sourced from one animal with behavioral signs of SCI-NP. A clear reduction in nociceptive responsiveness is evident immediately upon ISMS onset in this animal.

All surgical and electrophysiological procedures, including the spinal contusion injury, are consistent with those described previously (Bandres et al., 2021, 2023a; Bandres et al., 2022; McPherson and Bandres, 2021; McPherson et al., 2015). Briefly, however, a terminal electrophysiological experiment was conducted in all rats. For rats with SCI, the procedure occurred 6-8 weeks after SCI, at which point spontaneous recovery of motor function had plateaued and the neuropathic pain state was established (if it manifested).

We recorded extracellular neural activity throughout the dorso-ventral extent of the L5 spinal gray matter using a 32-channel microelectrode array. While recording, we induced natural nociceptive cutaneous feedback before, during, and after 30-minute epochs of sub-motor threshold ISMS delivered to the L5 spinal motor pools. Sensory feedback was induced by application of graded pressure to the most sensitive receptive field of the glabrous skin of the hindpaw ipsilateral to the implanted microelectrode array.

Trials had the following sequence: (a) 1 min of spontaneous neural transmission (i.e., no probing of the periphery); (b) 3-5 min of noxious mechanical stimulation of the receptive field: a ∼1-2 s pinch of the most sensitive region of the receptive field every 30 s; (c) 30 min of motor-targeted ISMS coupled with noxious mechanical stimulation of the receptive field; (d) 3-5 min of noxious mechanical stimulation of the receptive field without ISMS (as above, in b); and (e) 1 min of spontaneous transmission. The 30 min ISMS duration was selected to match that used previously to study the modulatory actions of motor-targeted ISMS on nociceptive specific and wide dynamic range neurons in neurologically intact rats and rats with chronic sensorimotor impairments following SCI (Bandres et al., 2023a; Bandres et al., 2022).

### Electrophysiology

We implanted a single microelectrode array (MEA) into the spinal cord at the L5 dorsal root entry zone (**Fig. 1A**). Each MEA contained two parallel shanks, and both shanks contained 16 discrete, vertically aligned electrodes (area: 177 μm^2^; inter-electrode spacing: 100 μm; NeuroNexus Inc., A2×16). The MEAs were electrodeposited with activated platinum-iridium to lower their impedance and increase their charge capacity (impedance: 4-10 KΩ; Platinum Group Coatings, Inc.). MEAs were also coated with 1,1’-Dioctadecyl-3,3,3’,3’-tetramethylindocarbocyanine perchlorate (Sigma-Aldrich, Inc.) to aid postmortem histological localization of the electrode tracks.

The MEA was lowered with a custom 4-axis, sub-micron resolution powered micromanipulator (Siskiyou, Inc.) until the bottom-most electrodes were located within the L5 dorsal roots. We then probed the glabrous skin of the ipsilateral hindpaw while monitoring dorsal root potentials. If dorsal root potentials were clearly correlated with periods of cutaneous feedback, we began to insert the MEA into the spinal cord. However, if dorsal root potentials were either not evident or if they were correlated with an inappropriate dermatome, we repositioned the MEA and repeated the mapping.

We then slowly advanced the MEA into the spinal cord, pausing every ∼25-50 μm to reduce shear and planar stress on the neural tissue. When the deepest electrodes reached the superficial border of the deep dorsal horn (∼400-500 μm deep), we again mapped the L5 dermatome while monitoring neural transmission in real-time. If multi-unit neural activity was clearly correlated with the desired receptive field, insertion continued; if the receptive field had shifted, the MEA was withdrawn and repositioned. When fully inserted, the ventral-most electrodes were positioned ∼1600-1800+ μm deep to the surface of the spinal cord, in the motor pools of the ventral horn. The MEA was not moved after full implantation.

ISMS was delivered as a series of discrete, cathode-leading pulses at 7 Hz, 200 μs/phase, 0 s inter-phase interval, and 90% of resting motor threshold (∼3-12 μA/phase). These ISMS parameters are consistent with values that induced functionally meaningful enhancements of spinal motor output below a chronic SCI (McPherson et al., 2015) and with our previous studies of the modulatory actions of motor-targeted ISMS on single-unit responsiveness to nociceptive sensory feedback (Bandres et al., 2023a; Bandres et al., 2022).

### Data processing and statistical analyses

All analyses were based upon features extracted from patterns of multi-unit neural transmission during nociceptive sensory feedback before, during, and after ISMS. Multiunit neural data (μV) from each channel was bandpass filtered between 750 – 15,000 Hz, detrended, rectified, and enveloped with a 250 ms moving root-mean-square function. For each pinch of the hindpaw, we extracted the voltage corresponding to the 95^th^ percentile per channel (**Fig. 1B**, red dots on left panel). The median per-channel 95^th^ percentile for each trial segment (i.e., pre-ISMS, during-ISMS, post-ISMS) was used to determine the extent to which that channel’s responsiveness to nociceptive feedback was potentiated, depressed, or unchanged during and following ISMS, respectively.

Electrode channels were also organized by anatomical region. These regions were defined as follows: superficial dorsal horn (sDH), associated with nociceptive transmission; deep dorsal horn (dDH), associated with nociceptive and non-nociceptive sensory transmission; intermediate gray matter (IG), associated with sensorimotor integration; and ventral horn (VH), most closely associated with motor-related transmission.

Statistical comparisons were based on repeated measures ANOVAs. Factors included anatomical region, number of channels potentiated or depressed at a given timepoint (relative to baseline, pre-ISMS discharge rates), rat, and an interaction term of anatomical region by number of channels potentiated vs. depressed. Post-hoc analyses compared the number of channels potentiated or depressed across anatomical regions; multiple comparisons were corrected using the method of Šídák. Comparisons were considered significant if the corrected *p* value was ≤ 0.05. Descriptive statistics presented in the text and figures display mean ± s.e.m. unless otherwise noted. Statistical analyses were performed using GraphPad Prism software (v9).

Animals were not randomized *per se*, although it was not possible to know *a priori* which animals would develop SCI-NP following injury. Behavioral analyses were conducted by an experimentalist not involved in terminal electrophysiological experiments, and it was not possible to determine during a terminal electrophysiological experiment whether a given animal did or did not exhibit behavioral signs of SCI-NP.

A power analysis was performed *a priori* (G*Power 3.1) using data extracted from a recent report detailing the impact of motor-targeted ISMS on discharge rates of wide dynamic range (WDR) and nociceptive specific (NS) neurons neurologically intact rats.(Bandres et al., 2022) Those data allowed estimation of an effect size for ISMS-associated depression of population discharge rates during nociceptive transmission. Achieved effect size in that study for repeated measures ANOVAs with the same independent and dependent variables as those used here was estimated to be 0.48, considered a medium effect by standard conventions. Using that effect size, α = 0.05, power (1-β) = 0.9, and a moderate correlation among repeated measures (0.6; range of 0-1, where the repeated measures are the mean discharge rate per rat across timepoints), 7 animals are required per group.

## RESULTS

It has previously been shown, both in neurologically intact rats and in a mixed cohort of rats with and without SCI-NP, that sub-motor threshold ISMS delivered in the motor pools depressed the majority of identified NS and WDR neurons’ responses to nociceptive transmission (Bandres et al., 2023a; Bandres et al., 2022). Left unresolved, however, has been the question of whether these single-unit responses are reflective of overall population-level behavior. Indeed, the prevailing hypothesis remains that deposition of electrical current into the spinal cord, whether intraspinally, epidurally, or transcutaneously, will have a net excitatory effect.

We first investigated population-level spinal responsiveness to nociceptive transmission in neurologically intact rats. Pooling across all regions of the spinal gray matter, 4/6 rats exhibited net depression of nociceptive responsiveness during and after ISMS compared to pre-ISMS baseline levels; two rats exhibited a net increase in nociceptive responsiveness during and after ISMS. This difference drove a significant main effect of the number of electrode channels potentiated vs. depressed both during and following stimulation (**Fig. 2A**; during ISMS: 33.51% of electrodes potentiated vs. 66.49% depressed, *p* = 0.049; following ISMS: 33.68% of electrodes potentiated vs. 66.32% depressed, *p* = 0.032).

**Figure 2.**
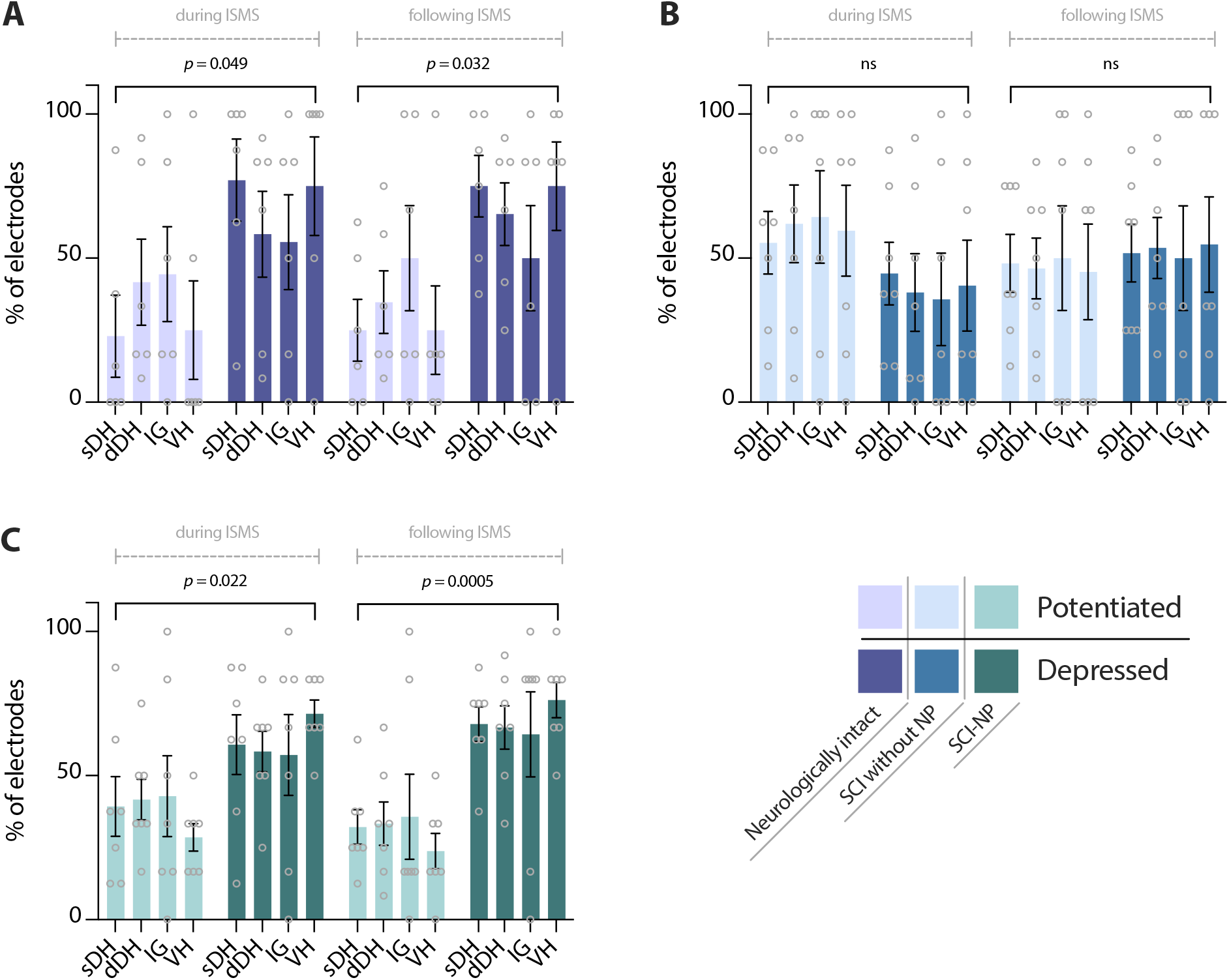
Motor-targeted ISMS preferentially reduces nociceptive responsiveness in rats with SCI-NP compared to rats that lack behavioral signs of SCI-NP. All subplots depict the percentage of electrodes exhibiting net potentiation (lighter shades) or depression (darker shades) during and following motor-targeted ISMS relative to pre-ISMS baseline levels. The percentages of electrodes potentiated vs. depressed are also presented relative to anatomical region in the spinal gray matter: sDH – superficial dorsal horn; dDH – deep dorsal horn; IG – intermediate gray; VH – ventral horn. (**A**) Neurologically intact animals (purple); (**B**) animals with chronic SCI that lack behavioral signs of SCI-NP (blue); (**C**) animals with behavioral signs of chronic SCI-NP (green). Note the increased percentage of depressed electrode channels relative to potentiated electrode channels in rats with SCI-NP compared to rats with SCI that lacked behavioral signs of SCI-NP.

We then subdivided the L5 spinal segment into regions commonly associated with distinct functional networks. This operation allowed us to determine if the overall depressive effect observed across electrodes was driven by preferential changes in certain regions. However, we found neither a significant main effect of region nor a significant interaction between region and the balance of depression vs. potentiation, either during or following ISMS. This finding implies that the modulatory capacity of ISMS is not modified by region.

We next asked if and how motor-targeted ISMS modulates population-level responsiveness to nociceptive sensory feedback in rats with chronic SCI. Although disinhibition of dorsal horn neurons is a feature common to sensorimotor incomplete SCI regardless of the presence or absence of SCI-NP, it is generally assumed that dorsal horn hyperexcitability is exaggerated further in rats with SCI-NP. Thus, we divided the overall SCI cohort into a subgroup of rats without behavioral signs of SCI-NP and a subgroup of rats *with* behavioral signs consistent with SCI-NP. Prior to ISMS, we found no significant differences between the mean amplitude of nociceptive responses (across electrodes) between neurologically intact animals, animals with SCI that lacked behavioral signs of SCI-NP, and animals with SCI-NP (*p* = 0.85).

Pooling across all regions of the spinal gray matter, 4/7 rats in the SCI cohort without SCI-NP exhibited net potentiation during and following stimulation and 3/7 exhibited net depression.

Although 60.27% of electrodes were potentiated across this cohort during ISMS compared to 39.73% depressed, the variability of responses rendered insignificant the main effect of number of channels potentiated vs. depressed (**Fig. 2B**, left; *p* = 0.16). Although we did observe a significant main effect of anatomical region (*p* = 0.01), there was not a significant interaction between anatomical region and the number of channels potentiated vs. depressed during ISMS (*p* = 0.97). Following stimulation, 47.47% of electrodes were potentiated compared to 52.53% depressed. This difference was not statistically significant (**Fig. 2B**, right; *p* = 0.73), and, expectedly, there was no interaction between anatomical region and the number of channels potentiated vs. depressed following ISMS (*p* > 0.99).

We next investigated the impact of ISMS on rats with SCI-NP. Given that ISMS did not reduce spinal responsiveness to nociceptive feedback in rats with SCI that lacked behavioral signs of SCI-NP – and qualitatively appeared to drive a modest (albeit statistically insignificant) facilitatory effect during stimulation – we predicted that ISMS would cause further potentiation of spinal responsiveness to nociceptive feedback in rats with SCI-NP. Surprisingly, however, this prediction was not supported by the empirical data. Six of 7 animals with SCI-NP exhibited a net *decrease* in nociceptive responsiveness during ISMS and all 7 animals exhibited a net decrease following ISMS. Nociceptive responses on 38.1% of electrodes were potentiated during ISMS across rats and regions during stimulation compared to 61.90% of electrodes depressed. This difference led to a significant main effect of the number of channels potentiated vs. depressed (**Fig. 2C**, left; *p* = 0.022). The main effect of region and the interaction of region and the number of channels potentiated vs. depressed were both insignificant. Following stimulation, 31.25% of electrodes were potentiated compared to 68.75% depressed, driving a significant main effect of the number of channels potentiated vs. depressed (**Fig. 2C**, right; *p* = 0.0005). Neither the main effect of region nor the interaction of region and the number of channels potentiated vs. depressed were significant.

## DISCUSSION

The primary finding of this study is that ISMS parameterized to enhance voluntary motor output after SCI reduces population-level spinal responsiveness to nociceptive sensory feedback during and following stimulation, both in neurologically intact rats and in rats with chronic SCI-NP. In contrast, motor-targeted ISMS did not robustly modulate nociceptive responsiveness in rats with chronic SCI that lacked behavioral signs of SCI-NP.

The most unexpected finding of this study was that motor-targeted ISMS reduced population-level nociceptive responsiveness in rats with SCI-NP but not in rats with SCI that lacked SCI-NP. The near uniformity of the depressive effect in rats with SCI-NP was also striking, and further distinguished the modulatory actions of ISMS on this cohort relative to the others. To the extent that ISMS was expected to modulate population-level nociceptive responsiveness at all after SCI, the most logical prediction was that the modulatory actions would be more robust in animals without SCI-NP. This prediction derives from previously published reports of the modulatory actions of motor-targeted ISMS on individual NS and WDR neurons in neurologically intact rats, rats with SCI-NP, and rats with SCI that lacked behavioral signs of SCI-NP (Bandres et al., 2023a; Bandres et al., 2022). It also derives from the presumably lower degree of dorsal horn hyperexcitability in rats that lack behavioral signs of SCI-NP compared those with an established pain state (Bedi et al., 2010; Carlton et al., 2009; Chen et al., 2022; Gwak and Hulsebosch, 2011; Nees et al., 2017). Yet, not only was the depressive effect more consistent in rats with SCI-NP than either of the other two cohorts, the magnitude of the depressive effect, as measured by the average percentage of electrode channels with potentiated vs. depressed responsiveness, was effectively the same in rats with SCI-NP and neurologically intact rats.

It was also surprising that the modulatory actions (or lack thereof) did not vary systematically with anatomical region in any of the cohorts. In this context, anatomical region can be considered a general proxy for network function (e.g., sDH being nociceptive-dominant; VH being motor-dominant). Anatomical region also has implications for the specific mixture of neurons composing the multi-unit spike trains accessed on each electrode channel. Presumably, some neuron types would be more responsive to nociceptive feedback than others. For example, channels composed of a greater proportion of NS and/or WDR neurons would be expected to be particularly sensitive to the modulatory actions of ISMS, given that these neurons naturally process nociceptive feedback. Considering that NS and WDR neurons are overwhelmingly located in the superficial and deep dorsal horns, one might have expected those regions to respond differently than the motor-dominant ventral horn. However, the magnitude of depression or potentiation was remarkably consistent across regions within each cohort.

### Potential mechanisms: pre-ISMS discharge rates

What could explain the primary finding that motor-targeted ISMS differentially modulated nociceptive responsiveness in rats with SCI-NP vs. those with SCI that lack SCI-NP? One explanation could relate to differences between the net level of dorsal horn excitability across the cohorts. Presumably, pathological overactivity in the dorsal horn of rats with SCI-NP is greater than that exhibited by rats without SCI-NP, which itself is elevated relative to neurologically intact animals. Canonical features of dorsal horn hyperexcitability include increased spontaneous and evoked discharge rates, lower recruitment thresholds for spinal neurons, and an increased number of spontaneously active neurons (Carlton, 2014; Gwak and Hulsebosch, 2011). Thus, the exaggerated firing dynamics of rats with SCI-NP would, at least mathematically, lead to a broader range over which spinal responsiveness to nociceptive feedback could be modulated. In this scenario, even if the intrinsic modulatory capacity of neurons was indistinguishable across cohorts, the effects of motor-targeted ISMS could paradoxically appear more robust in rats with SCI-NP.

Results from the cohort of neurologically intact rats argue against this interpretation, however. The average magnitude of evoked responses to nociceptive feedback before ISMS in these animals was indistinguishable from that of rats with SCI-NP and rats with SCI that lacked signs of SCI-NP. Yet, ISMS promoted a greater depressive effect on nociceptive transmission in neurologically intact animals than rats without SCI-NP, both in terms of the proportion of animals exhibiting net depression and in the percentage of potentiated vs. depressed electrodes. Thus, it appears that dorsal horn hyperexcitability – at least manifest as increased population-level evoked response amplitude – is not requisite to observe the depressive effects of motor-targeted ISMS.

If not a byproduct of increased baseline responsiveness, what else could underlie the differential effects of motor-targeted ISMS on rats with SCI-NP vs. those without? Given that the spinal contusion parameters and location were consistent across cohorts, it is unlikely that the effect is attributable to a systematic difference in the amount of descending neuromodulatory drive from the brainstem to spinal segments below the lesion. Thus, the differential effects presumably reflect distinct features of the modulated spinal segments themselves. Below, we nominate three potential mechanisms that could operate in isolation or concurrently to contribute the observed results. The veracity of these explanations could not be tested directly by the experiments detailed in this manuscript, and thus we urge caution in their interpretation.

### Potential mechanisms: intrinsic active membrane dynamics

First, the depressive effect in rats with SCI-NP could reflect the influence of strong of plateau potentials in many of the modulated neurons. In physiological conditions, an adequate level of neuromodulatory drive is required to instantiate the persistent inward currents (PIC) that support a plateau potential (Heckman et al., 2008; Morisset and Nagy, 1998). However, in pathophysiological conditions such as SCI, neuromodulatory drive is reduced or absent. In this case, active membrane properties undergo adaptive changes to preserve intrinsic cellular excitability, often resulting in maladaptively increased responsiveness to sensory feedback (and other excitatory synaptic inputs) due to denervation supersensitivity and/or constitutive activity of the metabotropic receptors that facilitate PICs (Fouad et al., 2010; Hains et al., 2003; Husch et al., 2012; Rank et al., 2007; Tysseling et al., 2017). For example, constitutive activity of dendritic serotonin receptors on motoneurons below an SCI is a key component of muscle spasms (D’Amico et al., 2013; Murray et al., 2010), and exaggerated plateau potentials in WDR projection neurons below an SCI are thought to be a key feature of SCI-NP (Derjean et al., 2003).

Interestingly, the PICs that support plateau potentials are exquisitely sensitive to synaptic inhibition (Bui et al., 2008; Hultborn et al., 2003; Kuo et al., 2003); in fact, inhibitory post-synaptic potentials are themselves amplified when the post-synaptic cell is under the influence of a strong PIC (Bui et al., 2008). If the depressive effect of ISMS is mediated at least in part by direct synaptic inhibition, then engagement of the same inhibitory networks in rats with SCI-NP and those without could theoretically lead to more pronounced depressive effects in rats with SCI-NP due to an increased influence of PICs and plateau potentials.

### Potential mechanisms: discharge state

Second, it is possible that ISMS promoted different neuronal discharge states across animals with SCI-NP and those without. WDR neurons have the capacity to manifest at least 3 discharge states: tonic firing (i.e., effectively tracking synaptic input 1-to-1), plateau potentials, and intrinsic bursting (Derjean et al., 2003), for example; motoneurons and other interneurons can likewise inhabit different firing states (Bandres et al., 2021; Dougherty and Chen, 2016; Lucas-Romero et al., 2022; Lucas-Romero et al., 2018; Steedman and Zachary, 1990; Stein et al., 2005). Each of these states has implications for the fidelity of information transfer through the cell. For example, tonic firing results in relatively faithful transmission of the input, whereas plateau potentials tend to amplify the input and intrinsic busting can exert either a facilitatory or depressive effect depending on the specific tuning of the inter-spike intervals relative to the membrane dynamics of the post-synaptic cell (Derjean et al., 2003; Izhikevich et al., 2003; Zeldenrust et al., 2018). Firing mode can switch dynamically depending on the balance of neuromodulation and synaptic inhibition.

In WDR neurons, endogenous bursting is associated with reduced transmission of afferent input (Derjean et al., 2003). Thus, if ISMS were to more readily induce or enhance bursting in rats with SCI-NP than rats without, the net effect at the population level would be depressed responsiveness to nociceptive transmission. It should be moted, however, that stable existence of WDR neurons in a bursting state appears to require activation of specific metabotropic glutamate receptors coupled with inactivation of GABA_B_ receptors (Derjean et al., 2003). Given that the neuromodulatory state of the spinal cord is likely to be somewhat similar across rats with SCI-NP and those without due to the conserved spinal contusion parameters, the role of ISMS in this scenario would presumably be to inhibit GABA_B_ receptors. It is not clear how such an effect would be realized, although it may involve actions on presynaptic GABA receptors rather than direct antagonism of GABA_B_ receptors on WDR dendrites.

### Potential mechanisms: latent inhibitory network

Finally, it is possible that ISMS engaged a latent inhibitory network in rats with SCI-NP that remained dormant in rats without SCI-NP. The existence of such a network is purely speculative, and neither its architecture nor its functions can be inferred from this study. In the context of intrinsic active membrane dynamics, however, two aspects of such a network are worth noting. First, if such a network were to promote presynaptic release of GABA in the vicinity of GABA_B_ receptors on WDR dendrites, it could reduce nociceptive responsiveness not by inhibiting the neuron *per se*, but rather by forcing a change in firing state from plateau to bursting or tonic.

And second, it has been demonstrated that sources of synaptic inhibition more effectively reduce discharge rates in neurons expressing PICs when those neurons are recruited by sources of synaptic excitation rather than by direct current injection at or near the soma (e.g., from a microelectrode) (Hultborn et al., 2003). The ISMS protocol used here may have mimicked this situation. Indeed, it is well-established that ISMS recruits low threshold interneurons and fibers of passage at sub-motor threshold intensities such as that used here (Gaunt et al., 2006; Gustafsson and Jankowska, 1976; Jankowska and Roberts, 1972). ISMS was also delivered in the motor pools, distant from the majority of classically defined spinal nociceptive pathways.

Considering that charge density decays as the square of radial distance from the electrode tip (Tehovnik, 1996), these aspects of the protocol together support the conclusion that the modulatory actions of motor-targeted ISMS on nociceptive transmission were unlikely to have resulted from direct current spread. Rather, they support the conclusion that the effects were driven by synaptic actions. As aforementioned, if PICs are more prevalent in animals with SCI-NP than animals without behavioral signs of SCI-NP, then motor-targeted ISMS may have been uniquely capable of capitalizing on this inhibitory amplification in the SCI-NP cohort.

### Limitations

There are several important considerations to reiterate about this study. For example, all experiments were conducted in anesthetized rats due to the invasive nature of the *in vivo* electrophysiological studies. And although we and others have demonstrated that urethane preserves the excitability of spinal nociceptive networks while exerting a relatively modest effect on GABA-ergic and excitatory amino acid transmission compared to other anesthetics (Bandres et al., 2021, 2023a; Bandres et al., 2022; Dalo and Hackman, 2013; Hara and Harris, 2002; Maggi and Meli, 1986a, b, c), it should not be assumed that the modulatory actions observed here will persist unchanged during awake, behaving conditions. This study was also unable to determine the extent to which neural transmission on a given electrode channel was dominated by output neurons vs. local, segmental interneurons, further precluding a direct mapping of the observed effects onto the behavioral experience of pain.

### Conclusion

This study demonstrates that the anti-nociceptive effects of ISMS intended to enhance voluntary motor output generalize to patterns of population-level neural transmission. In so doing, it motivates future work to optimize these multi-modal effects. Such work could involve investigating different stimulation parameters (e.g., stimulation frequency, pattern, location), electrode montages, and/or closed-loop configurations. This study also demonstrated, unexpectedly, that the anti-nociceptive effects of motor-targeted ISMS are most robust in animals with SCI-NP. A similar finding in awake, behaving animals would represent an important translational milestone, suggesting that the multi-modal effects of motor-targeted ISMS may be most efficacious in the individuals most in need of such a therapy, i.e., people living with chronic motor impairments and neuropathic pain following SCI.

## ABBREVIATIONS

dDH: deep dorsal horn
IG: intermediate gray matter
ISMS: intraspinal microstimulation
L5: 5^th^ lumbar spinal segment
MEA: microelectrode array
NS: nociceptive specific neuron
PIC: persistent inward current
SCI: Spinal cord injury
SCI-NP: spinal cord injury-related neuropathic pain
sDH: superficial dorsal horn
VH: ventral horn
WDR: wide dynamic range neuron

